# Coevolution of species colonization rates controls food-chain length in spatially structured food webs

**DOI:** 10.1101/2022.10.11.511705

**Authors:** Vincent Calcagno, Patrice David, Philippe Jarne, François Massol

## Abstract

The complexity of food webs and how it depends on environmental variables is a long-standing question in ecology. Food-chain length depends on many factors, such as productive space, disturbance and spatial processes. It is not clear though how food-chain length should vary with adaptive evolutionary changes of the constitutive species. Assuming that each trophic level is subject to a competition-colonization trade-off, we model the adaptive evolution of their colonization rates and its consequences on equilibrium occupancies and food-chain length. When colonization rates are allowed to evolve, longer food chains can persist. Extinction, perturbation and habitat loss all affect the evolutionary-stable colonization rates, but trade-off strength (costs of dispersal) has a major role: weaker trade-offs yield longer chains. Our results suggest a strong link between within-trophic level competition, spatial occupancies and food-chain length. Although these eco-evo dynamics partly alleviate the constraint on food-chain length in metacommunities, we show it is no magic bullet: the highest, most vulnerable, trophic levels are also those that least benefit from evolution. Our model generates qualitative predictions regarding how trait evolution affects the response of communities to disturbance and habitat loss. This highlights the importance of eco-evolutionary dynamics at metacommunity level in determining food-chain length.

## Introduction

An important puzzle in ecology is how food web topology, and in particular food-chain length, is determined (Hutchinson, 1959, Pimm, 1982, May, 1983, Cohen and Briand, 1984, Stenseth, 1985, Cohen and Newman, 1988, Williams and Martinez, 2000, 2004). Food-chain length is a measure of the number of feeding links between resources and top predators (*e.g.* Sabo et al., 2009). Ecological theory has long tried to understand what limits this length (Hutchinson, 1959, May, 1972, Hastings and Conrad, 1979, Pimm, 1982, Menge and Sutherland, 1987). For instance, the energetic constraint hypothesis (Hutchinson, 1959) invokes imperfect transfers of energy and resources along food chains, the dynamics constraint hypothesis (May, 1972, Pimm and Lawton, 1977) argues that long food chains are more vulnerable to perturbation than short ones, and the community area hypothesis combines the diversity-area relationship obtained by the theory of island biogeography (MacArthur and Wilson, 1967) with the link scaling law (Cohen and Briand, 1984) to predict a concave increase in food-chain length with habitat area (Cohen and Newman, 1991). Recent empirical studies have identified three major determinants of food-chain length: productive space (*i.e.* ecosystem size × productivity), disturbance, and ecosystem size (Post, 2002). While confirming the roles of resource limitation and perturbation, these results argue against single explanations, and also stress the need to incorporate space in theoretical models. Indeed, despite ample evidence that food-chain length correlates with habitat area or ecosystem size (Schoener, 1989, Cohen and Newman, 1991, Post et al., 2000, Takimoto et al., 2008), spatial processes are still understudied in theoretical models of food webs (Holt, 2002, Amarasekare, 2008). Calcagno et al. (2011) extended the model of Holt (2002) to provide predictions under a range of conditions encompassing top-down effects of any sorts and habitat selection effects, while Pillai et al. (2011) studied the effect of colonization rates on the complexity of food webs at the metacommunity level. More recently, Wang et al. (2021) reanalyzed a similar ecological model of spatial food chains in order to study how metacommunity dynamics should influence maximum food-chain length. They rendered the model spatially realistic and provided some empirical support for its predictions in a butterfly-parasitoid system.

The spatial structure of populations, on the other hand, has inspired one of the most prolific research lines in evolutionary biology, namely the evolution of dispersal (for reviews, Bowler and Benton, 2005; Ronce, 2007; Duputié and Massol, 2013). Indeed, there has been an early realization that spatial structure, beyond its population dynamical consequences, also imposed strong selection pressures, and that the key parameter in this context, the colonization, dispersal, or migration rate, could be shaped by evolutionary adaptation. A range of selective forces should act on the dispersal rate, and determine its evolutionarily stable (ES) value, depending among others on patch size, connectivity, spatio-temporal environmental variability and dispersal costs. In the case of metapopulation occupancy models, following the seminal work of Levins (1969), Slatkin (1974), Hastings (1980), Hanski (1983), Tilman (1994) and others, one can translate a purely ecological statement of the problem (“what is the occupancy of competing species when they obey a competition-colonization trade-off?”, Calcagno et al. 2006) into an evolutionary question (“how should this competition-colonization trade-off drive the evolution of colonization rates of coexisting species?”, Aubrée et al. 2020), and thence investigate the eco-evolutionary dynamics of adaptation of colonization rates in a metacommunity.

The purpose of this article is to connect these two important aspects of spatially structured communities, namely the ecological limit they impose on food-chain length on the one hand, and the evolutionary dynamics of dispersal on the other hand, and to determine how they could interact and influence one another. In Calcagno et al. (2011), two distinct constraints on food-chain length arising from metacommunity structure were identified. First, finite colonization rates limit predator occupancy to a subset of prey-occupied sites. Second, extinction rates accumulate along food chains. Both processes thus concur to decrease maximal and average food-chain length in metacommunities. However, in an eco-evolutionary dynamics framework, the evolution of colonization rates, in response to competition within trophic levels, will affect the first constraint (by either increasing or decreasing colonization rates), and hence will likely change trophic level occupancies and food-chain length. Here, we investigate this question theoretically by plugging an adaptive dynamics-based model of colonization evolution into the food-chain metacommunity model of Calcagno et al. (2011). This model allows us to assess how habitat availability and extinction and perturbation rates select for colonization rates, and how this selection changes prediction between a metacommunity that evolves and one that does not. The parameters of the competition-colonization trade-off provide us with another lever quantifying the strength of intra-trophic level competition. Ultimately, this model allows us to predict how food-chain length changes with environmental variables in response to selection on colonization rates.

## Model and Methods

### Spatial food chain model

We use a model of spatial food chain identical to that in Calcagno et al. (2011), and very close to that of Wang et al. (2021). It is best formulated in terms of the dynamics of 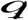, the fraction of patches that contain trophic level *i*, and thus all the lower trophic levels too, regardless of the presence/absence of upper trophic levels (Fig. 1). These dynamics can be expressed in a way similar to Levins’ classical metapopulation model:

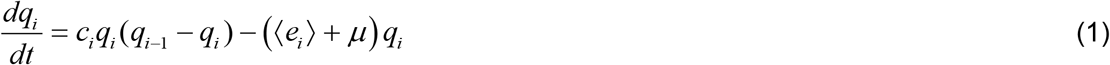

where for the first trophic level (*i*=1), *q*_0_ = *h* is the proportion of habitat (patches) available (level 0 is therefore not counted in food-chain length). As a simple example, a fully developed set of equations is described in Supplementary Material for a two-level trophic chain.

**Figure 1.**
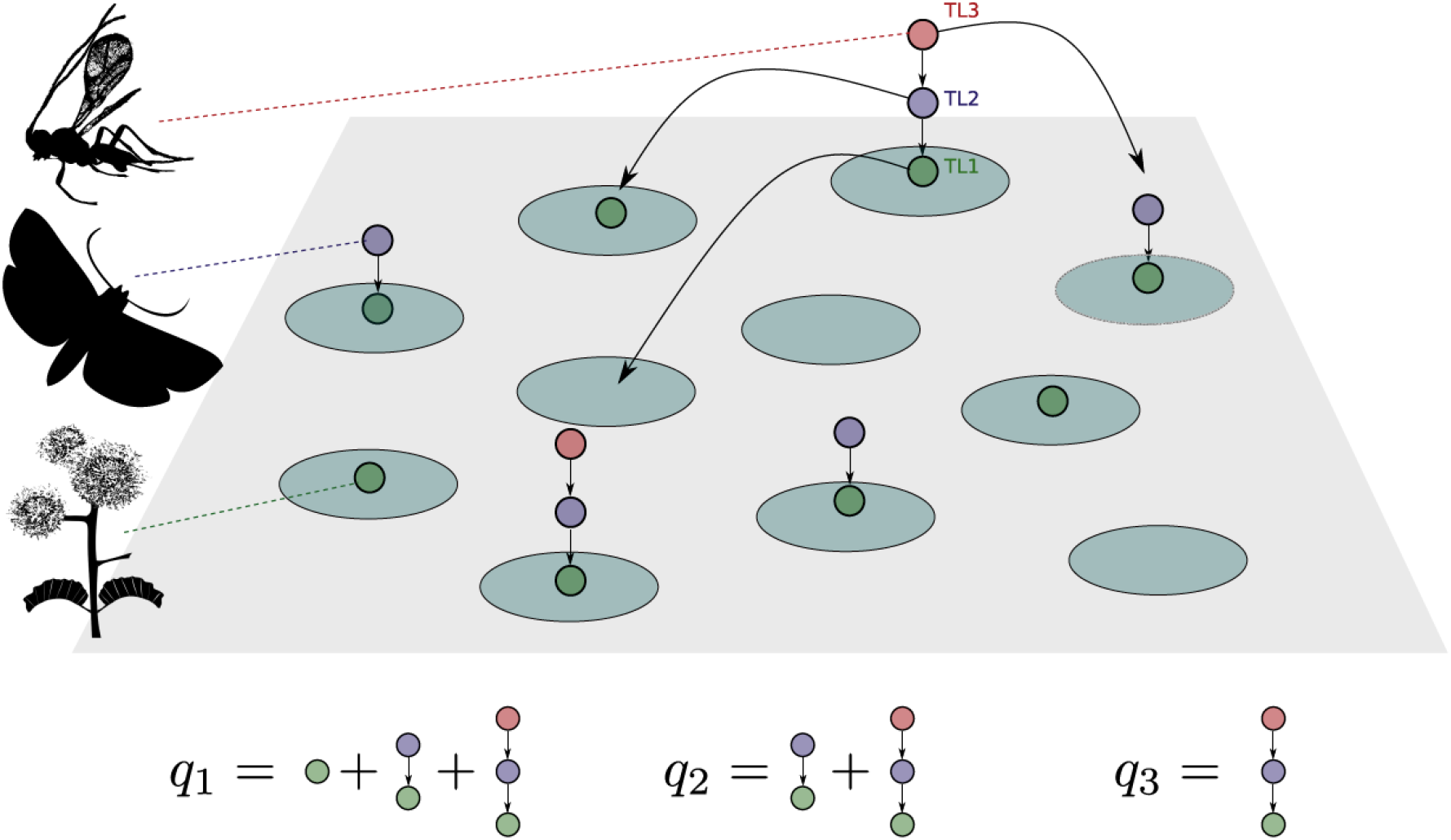
A graphical illustration of the spatial food chain model. The habitat consists of many interconnected patches. Each patch can be colonized by one or more trophic levels. The first trophic level can colonize any empty available habitat patch, whereas the second trophic level can only colonize patches where the first trophic level is found, etc. Each trophic level can disappear from a patch at some basal extinction rate *e*, and local extinction of one trophic level causes the local extinction of all upper trophic levels. Patches can also undergo catastrophic perturbation events, at rate *μ*, causing all trophic levels to be removed at once. The dynamical variables of interest in eq. (1) are the *q_i_*, *i.e.* the proportion of patches occupied by the *i*-th trophic level (and possibly upper trophic levels too), as illustrated at the bottom of the figure.

This equation assumes that patches experience perturbation at rate *μ*, which causes all trophic levels to disappear simultaneously (perturbation-driven extinction). An additional fraction of extinctions, called the effective extinction rate 〈*e_i_*〉, is experienced independently by each trophic level, and incorporates a basal extinction rate *e*, potentially modified by bottom-up and top-down effects. Bottom-up effects reflect the fact that if a given trophic level goes extinct in a patch, all upper trophic levels must go extinct too. Top-down effects represent how the presence of upper trophic levels may increase the extinction rate of the focal trophic level (parameter *e_TD_*), for instance through a reduction of local population size. We assume that in the fraction of sites that contain the *i*+1 trophic level (= *q*_*i*+1_/*q*_*i*_) top-down effects increase the extinction rate by a constant. Applying this rule to all trophic levels, this yields 〈*e_i_*〉 to be expressed as *e* + (*i* − 1)(*e* + *e_TD_*) + *e_TD_q*_*i*+1_ / *q_i_* (see Calcagno et al. 2011; Wang et al. 2021).

### Evolutionary dynamics of colonization rate

The above model has been thoroughly studied for arbitrary, fixed, colonization rates of the different trophic levels (Calcagno et al. 2011). In this article, we further let colonization rate values evolve under the effect of natural selection (the basal extinction rates and the strength of top-down effects are fixed). Metapopulation structure is known to impose strong selection pressures on dispersal, and hence on the colonization rate (Duputié & Massol 2013). Dispersal is selected for by the presence of empty available patches (Comins et al. 1980), and, for low enough population sizes, kin competition. On the other hand, dispersal is counter-selected by competitive trade-offs, *i.e.* the fact that high dispersal rates come at the expense of local growth and/or competitive strength, which is the case when dispersal is costly (Supplementary Material). We here use a simple representation of these evolutionary dynamics, that can be derived from competition-colonization trade-offs (Calcagno et al. 2017), or alternatively from spatially structured population models (Supplementary Material). It incorporates the three selective forces listed above. The derivation yields the following equation for the dynamics of *c_i_*:

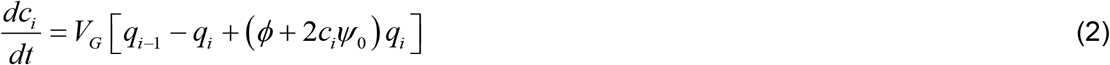

*V_G_* is a parameter representing the genetic variance in colonization (from a quantitative genetics perspective, Lande 1979; see Dieckmann & Law 1996 for an alternative interpretation). The other two evolutionary parameters describe the strength of the trade-off between local competitiveness and colonization (Supplementary Material). *ϕ* is the probability that immigrants replace residents when they have the same colonization rate (between 0 and 1), and *ψ*_0_ is the (negative) slope quantifying how much increasing colonization rate comes at the expense of decreased local competitiveness (*i.e.* there is a cost of dispersal). The first two terms in the brackets represent positive selection for dispersal caused by the presence of empty available patches (fraction *q*_*i*−1_ − *q_i_*), and the third term (*ϕ*+2*c_i_ψ*_0_)*q_i_* represents the usually negative selection on colonization rate caused by local competition. More specifically, the *ϕ* term represents the (weak) positive selection caused by kin competition (Supplementary Material), whereas the term involving *ψ*_0_ represents the counter-selection imposed by competitive trade-offs and costs of dispersal. As shown in Supplementary Material, eq. (2) admits a unique positive equilibrium in the cases of interest; this singular strategy is always convergence stable, and is an evolutionarily stable strategy (ESS) provided one condition is met by the trade-off function. The ESS is attained when the selective forces for and against colonization cancel out.

### Computing equilibria and maximum food-chain length

The dynamics of (1) can be used directly, with fixed colonization rates, to compute food-chain length in the absence of evolution (Calcagno et al. 2010; Wang et al. 2021). To introduce evolution, we study the coupled dynamics of eq. (1) and (2), using the common simplification that eq. (1) was at equilibrium, *i.e.* that metapopulation dynamics was faster than evolutionary dynamics. Given the nature of the ecological and evolutionary attractors, the assumption is not expected to have important consequences (Sanchez et al. 2011). For numerical examples with small numbers of trophic levels, we numerically integrated eq. (2) through time, keeping eq. (1) at equilibrium. For larger numbers of trophic levels and to compute maximum food-chain length, we assumed eq. (2) was at equilibrium (ESS), and implemented an iterative algorithm allowing to jointly solve eq. (1) and (2) (see Supplementary Material for details on the algorithm). Using this algorithm, we computed the maximum food-chain length, defined as the maximum number of trophic levels that can persist, where persistence is defined as having equilibrium occupancy greater than some small threshold level *ϵ*. In practice, we used 10^−3^, meaning that any trophic level should occupy at equilibrium at least one patch in one thousand in order to persist.

In all figures, we used the following reference parameter values, unless stated otherwise or a parameter was systematically varied: *h* = 1, *e* = 1, μ = 1, *ϕ* = 0.5, *ψ*_0_ = −0.1.

## Results

### The evolution of dispersal in a spatial food chain

Fig. 2a-b presents results for the simplest possible case with only two trophic levels: one prey and one predator. In the figure, the basal extinction rate is gradually increased, causing the occupancy of the two trophic levels to decline, eventually going extinct one after the other. This basically mimics a continuous degradation of the environment which becomes harsher and harsher in time, as expected for example if the taxa are challenged by climate change or anthropisation. In the absence of evolution, the figure was parameterized in order to replicate the results of Wang et al. (2021)’s figure 1, thus including top-down control of the predator. In that case, our mathematical model faithfully reproduces these spatially realistic simulation results. Occupancy decreases markedly for the higher trophic level, which disappears at rather low extinction rates. The lower trophic level can maintain itself at much larger extinction rates. Due to the absence of trait dynamics, he colonization ratio (*c/e*) curves remain identical, until the higher trophic level disappears.

**Figure 2.**
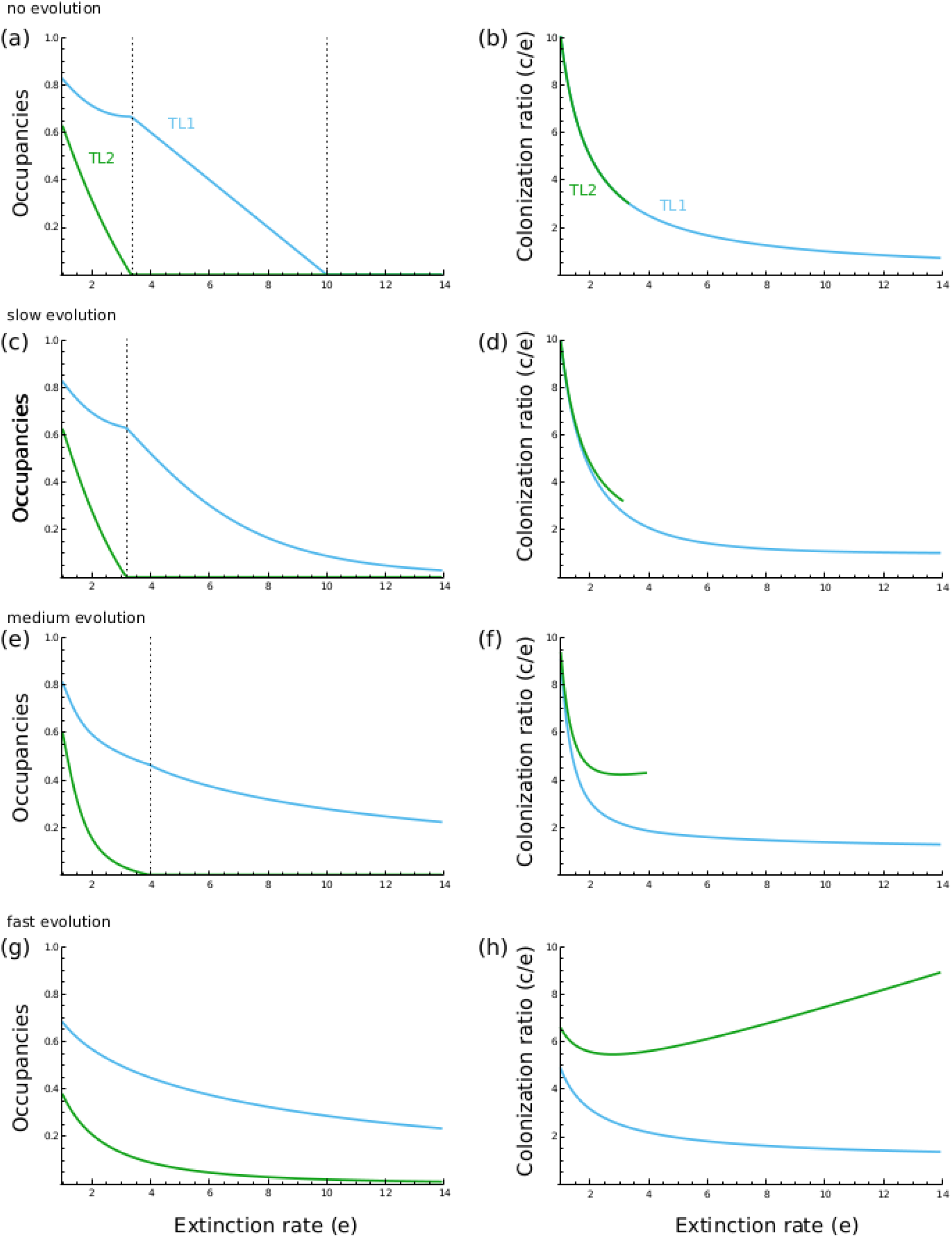
Increasing extinction rate in a system with two trophic levels. (**a-b**) No evolution. The extinction rate is gradually increased from its baseline value (*e*=1). The equilibrium occupancies of the prey (*blue*) and the predator (*green*) gradually decline until extinction (a), as the colonization ratios (*c/e*) decline with the increase in *e* (b). These results very closely reproduce Wang et al. (2021) fig. 1 results. (**c-d**) Introducing evolution at low speed relative to the change in *e* (*V_G_* = 0.1). (**e-f**) Introducing evolution at an intermediate speed relative to the change in *e* (*V_G_* = 1). (**g-h**) Introducing evolution at a large speed relative to the change in *e* (*V_G_* = 100). The two trophic levels initially had colonization rates *c* = *10*; without evolution they retained these values throughout, whereas with evolution the values gradually changed according to eq. (2). Other parameters were at default values, except *μ* = 0 and *e_TD_* = 2 *e*.

Fig. 2c-h shows the impact of evolution on these results. Important quantitative and qualitative differences emerge. If evolution is slow (Fig. 1c-d), *i.e.* if parameter *V_G_* is low in eq. (2), evolution has little impact, and does not significantly “delay” the extinction of the predator (*i.e.* allows it to survive at higher values of *e*). Despite the predator evolving higher colonization rate when approaching extinction, eventually outpacing the increase in extinction rate (*i.e.* the colonization ratio *c/e* increases), this is not sufficient to prevent extinction. Following the predator’s extinction however, the prey manages to persist despite the continuous increase in extinction rate, through a gradual increase in its colonization rate, and thus its colonization ratio *c/e* (Fig. 2d). Its occupancy declines, but the decline is mitigated by evolution, preventing extinction in the range of extinction rates considered.

When evolution is faster, the predator can evolve sufficiently to delay its extinction (Fig. 2e-f), and ultimately it also manages to escape extinction (albeit at low occupancy), just like the prey (Fig. 2g-h). Note that when the prey is not threatened with extinction, evolution in the prey can initially decrease its equilibrium occupancy, which is detrimental to predator persistence and counteracts the effects of evolution in the latter.

From eq. (2), it can be seen that as a species approaches extinction, *q_i_* → 0, *c_i_* always increases, acting to counteract extinction in all cases. However, comparing among trophic levels when they are on the verge of extinction (*q_i_* ≈ *ϵ*), one can deduce that (i) the positive selection pressure caused by available habitat is lower for higher trophic levels (since *q*_*i*−1_ decreases with *i*), and (ii) the negative selection pressure caused by local competition is stronger (since the colonization rate *c_i_* will generally be increasing with trophic position). Both effects make selection for larger colonization rates less efficient at higher trophic levels, providing a rationale for the results shown in Fig. 2.

Overall, the evolution of colonization rate is beneficial to species persistence and thus food-chain length. Higher trophic levels will be selected for larger colonization rates than lower trophic levels. However, while all trophic levels are selected for higher colonization when approaching extinction, the intensity of the effect increases with trophic level, putting a higher evolutionary challenge on higher trophic levels. Furthermore, evolution at low trophic levels, when they are not threatened by extinction, is often detrimental to the persistence of upper trophic levels. For these two reasons, the beneficial effects of evolution are not as effective as one might expect to promote food-chain length. Basically, a fast evolution of predators is more efficient at maintaining the integrity of the food chain than a fast evolution of their prey; fast evolution in prey becomes necessary to maintain the system only in conditions where the prey would be threatened even in the absence of predation.

Since results go in the same direction with fast and slow evolution and since assuming the ESS allows for much easier mathematical analysis and faster computing (Supplementary Material), in the following we will consider fast evolution only, and compare results with the “no evolution” scenario. Our results can be taken as giving an upper bound on the potential consequence of evolution on food-web dynamics, and capturing the two extreme possible situations, most actual situations probably falling somewhere in between them.

### Maximum food-chain length under habitat degradation

We considered two forms of habitat degradation: (i) an increase in extinction rate (as in the previous section), and (ii) habitat destruction (*i.e.* a decrease in the fraction of available patches *h*; see equation [1]).

As expected, increasing the extinction rate decreases the maximum food-chain length in the absence of evolution (Fig. 3a, see also Calcagno et al. 2011; Wang et al. 2021). In the presence of evolution, however, the decline is much reduced. This is caused by an increase in the colonization rate of the upper trophic levels, which are selected for high values, even though this does not suffice to overcome the increasing extinction rate and the reduced occupancy of lower trophic levels (Fig. 3b). Note that, with or without evolution, the decline of maximum food-chain length with extinction rate always has an overall convex shape (rapid loss of many trophic levels followed by gradual loss of the remaining levels as extinction rate increases).

**Figure 3.**
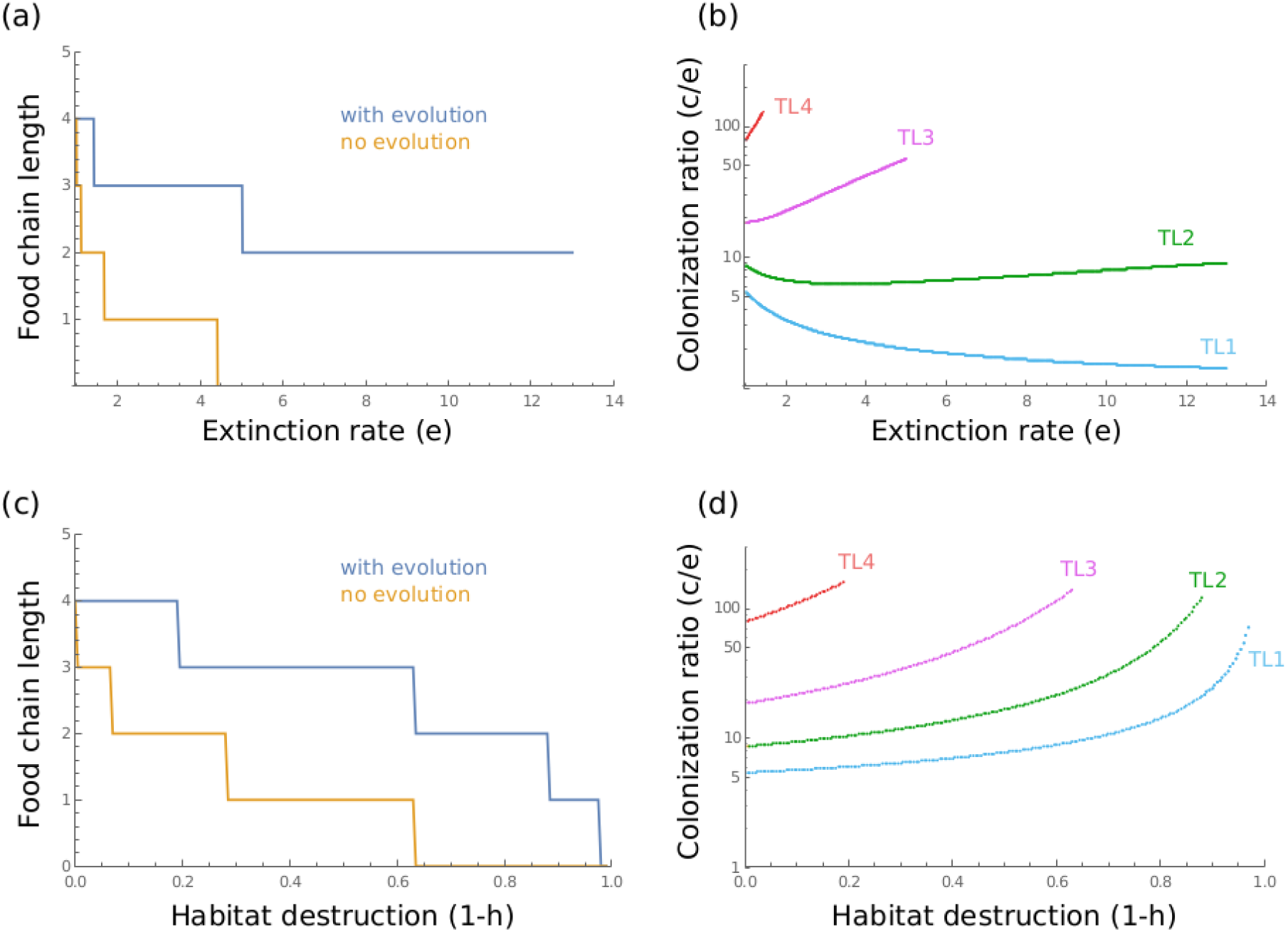
Maximum food-chain length under habitat deterioration. (**a-b**) The effect of increasing the extinction rate on maximum food-chain length (a), without evolution (*orange*) and with evolution (*blue*). In the case of evolution, the colonization ratio values at each trophic level are shown in (b). (**c-d**) Same as (a-b), except that the level of habitat destruction (1-*h*) was increased instead of *e*. Initially (*i.e.* for *e*=*0* or *h*=*1*), species have the same colonization rates in the two scenarios (with or without evolution), set at the corresponding ES values. These values do not change in the no evolution scenario, whereas they changed in the other scenario.

When destroying habitat patches (reducing *h*), the same tendency is observed: evolution allows longer food chains to persist, compared to the reference case with no evolution (Fig. 3c). Note, however, that all trophic levels jointly evolve higher colonization rates, even the lower ones (Fig. 3d). This is because decreasing *h* affects the first trophic level just as strongly as the others, unlike increasing the extinction rate, which has a disproportionately stronger effect on the higher trophic levels. We also remark a qualitative impact of evolution: the decline of food-chain length with habitat destruction follows a convex sequence in the absence of evolution (first many losses, then more gradual losses), as it does when manipulating extinction rate. However, with evolution, increasing habitat destruction causes a concave declining sequence of food-chain length, with two trophic levels (out of four) going extinct when habitat destruction attains values of 0.8-1.0 (Fig. 3c).

### The consequences of trade-off intensity

Increasing the intensity of the trade-off between colonization and local competitiveness (*i.e.* decreasing the basal competitiveness of immigrants *ϕ* and/or making the slope *ψ*_0_ more negative) increases the negative selection pressure on the colonization rates (eq. 2), and should thus select for lower values of the latter. This should in turn negatively affect maximum food-chain length. This is indeed what we find (Fig. 4a), except that parameter *ϕ* has very little impact. The latter result is consistent with Calcagno et al. (2017). If the trade-off slope is very shallow, colonization rates are free to evolve to quite high values, which allows much longer food chains to persist (Fig. 4b). Conversely, if the slope is very negative, selection favours low colonization values, especially at lower trophic levels, at the expense of population persistence and food-chain length (Fig. 4b). Costs of dispersal (trade-off intensity) are therefore a critical aspect for predicting the consequences of dispersal evolution for food-chain length.

**Figure 4.**
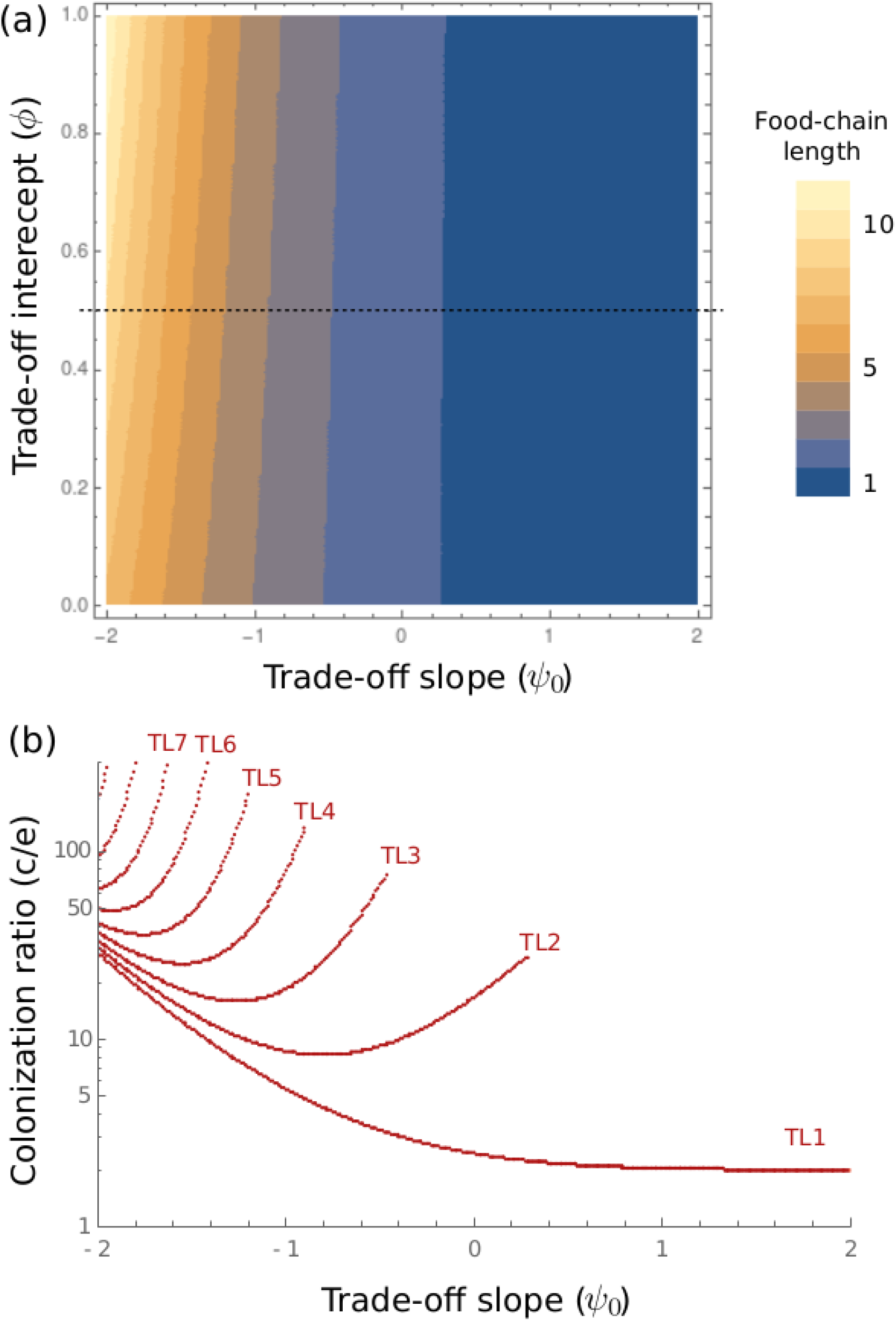
Consequences of varying the competitive trade-off intensity. (**a**) A contour plot of maximum food-chain length, with evolution, as a function of the two competitive trade-off parameters (*ϕ*; y-axis) and (*ψ*_0_; log scale, x-axis; −100 < *ψ*_0_ < −0.01). (**b**) For a fixed *ϕ* = 0.5 (corresponding to the dotted line in panel a), the colonization ratios of the different trophic levels as a function of *ψ*_0_.

### The relationship between food-chain length and the proportion of empty patches

As demonstrated by Calcagno et al. (2011) and Wang et al. (2021), in the absence of evolution, a negative correlation is expected between food-chain length and the fraction of empty patches. In other words, long food chains can be observed only if the total occupancy is high enough, as shown in Fig. 5. When varying the extinction rate *e*, we again find that evolution changes the pattern quantitatively, but not quantitatively (Fig. 5a-b). Things are different when the amount of habitat destruction varies: in this case, we observe concave negative relationships between food-chain length and the fraction of empty patches, but only with evolution. Without evolution, we observe the classic convex relationships observed otherwise or, at best, linear relationships (Fig. 5c-d). Such a qualitative difference in the shape of the relationships could be looked for in empirical or experimental data in order to infer the action of evolutionary dynamics. Interestingly, this difference is reminiscent of the one we have observed between food-chain length and habitat destruction (Fig. 3c-d). We checked that varying other parameters (specifically the perturbation rate *μ*) could not produce concave relationships, with or without evolution (Supplementary Fig. 2). It therefore seems that habitat destruction has a specific action that brings up a qualitative difference between evolved and unevolved food chains. Specifically, with evolution, relationships that are otherwise convex can be turned concave.

**Figure 5.**
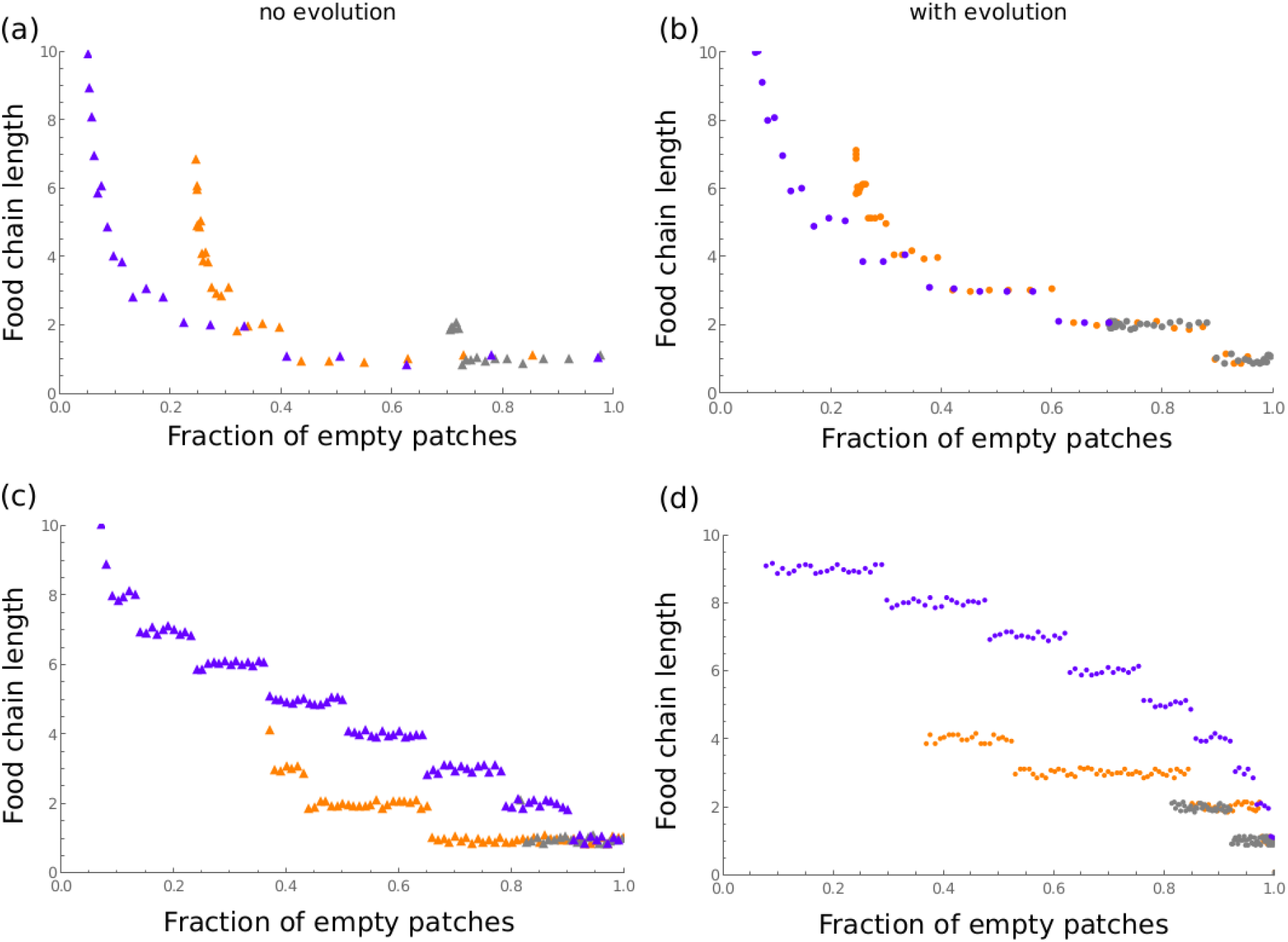
The relationship between food-chain length and the fraction of empty patches without evolution (left panels) and with evolution (right panels). (**a-b**) Each dot represents a different parameter value, where the extinction rate varies between 0.01 and 100, under three different values of *ψ*_0_. (**c-d**) Each dot represents a different parameter value, where the habitat destruction level 1-*h* varies between 0 and 1, under three values of *ψ*_0_. In all panels, the three values of *ψ*_0_ were −1 (gray dots), −0.1 (orange dots) and −0.01 (blue dots). As in Fig. 3, species had colonization rates set at the ES values for the initial parameter value, and these values remained constant in the no evolution scenarios, whereas they adapted in the evolution scenarios. Points were slightly jittered vertically for better readability.

## Discussion

As previous studies have shown, metacommunity dynamics impose an intrinsic constraint on food-chain length, and the persistence of long food-chains in spatially structured habitats thus requires specific conditions (Holt, 2002; Calcagno et al. 2011). Higher trophic levels are vulnerable to extinction from multiple processes, and, paraphrasing Robert May, they must arguably deploy “devious strategies” in order to persist. The most obvious strategy in this context is the adoption of systematically higher colonization rates by higher trophic levels, but there are also other strategies, such as behavioural ones (Calcagno et al. 2011).

As shown here, the evolutionary dynamics of colonization rates, *i.e.* the evolution of dispersal at individual trophic levels, is yet another way to alleviate the constraint on food-chain length. Natural selection does favour larger colonization rates at upper trophic levels and an evolutionary increase in colonization rate can, if fast enough, allow the persistence of longer food chains than when species traits are constant. The selection pressures imposed on colonization rates by trophic relationships and spatial structure cause evolutionary responses that can sustain the persistence of more trophic levels in the face of habitat degradation, such as increased disturbances or habitat destruction.

It would seem at first glance that such eco-evolutionary dynamics could be the ultimate devious strategy, by allowing the colonization rates of species to evolve larger values as needed. They could potentially alleviate any limit on food-chain length, as spatial persistence is ensured provided a sufficiently large colonization rate. This would be similar to evolutionary rescue, in another context and at a larger scale. However, our results show that things are not so simple. Indeed, when all trophic levels are allowed to evolve in such a way, two factors limit the efficiency of evolution in rescuing food-chain length. First, the lowest trophic levels, which are typically not facing the greatest danger of spatial extinction, experience little selection for increased colonization as their effective extinction rates are low. Quite the opposite, they can readily be selected for lower colonization rates, because of intraspecific competition or, equivalently, costs of dispersal. Such a negative selection pressure lowers their spatial occupancy, directly jeopardizing the persistence of the superior trophic levels. Second, the strength of selection for larger colonization rates typically declines, all else being equal, as trophic level increases. This is because positive selection declines as the fraction of hospitable patches decreases, while at the same time, the larger colonization rates that higher trophic levels must adopt increase the strength of selection against dispersal caused by intraspecific competition (see the gradient equation [2]). Combined, these two factors make eco-evolutionary dynamics at lower trophic levels potentially detrimental to maximum food-chain length, and not so effective at rescuing higher trophic levels. In short, even if very efficient, evolutionary dynamics is not the silver bullet to ensure the persistence of higher trophic levels. This is why, even with no particular constraint on the speed or extent of adaptive evolution, eco-evolutionary dynamics does not totally suppress the spatial constraint on maximum food-chain length. Even when it permits longer food-chains to persist, the higher trophic levels are typically kept at low spatial occupancies, thus vulnerable to environmental perturbations or other random events.

In the evolutionary context we have considered here, the question of understanding what devious strategies higher trophic levels must deploy to persist remains, though the range of available strategies is more precisely defined. Theoretically, factors that could favour the persistence of longer food-chains are: (i) higher trophic levels have faster evolution rates, or (ii) competition-colonization trade-off intensities are weaker at higher trophic levels. These two possibilities remain to be investigated empirically. We can nonetheless remark that one of the components affecting trade-off intensity is local population size (Supplementary Material). The larger the local population size, the less steep the trade-off, all else being equal. Since we would quite often expect higher trophic levels to have smaller local population sizes, this should make *ψ*_0_ more negative as trophic level goes up. If so, this would be yet another factor detrimental to food-chain length under eco-evolutionary dynamics. Regarding heterogeneity in evolution rates among trophic levels, the good news is that, if evolution proceeded more quickly in prey than predators or vice versa for all traits under selection, such a heterogeneity should leave a clear mark on the population dynamics series of both predator and prey (Yoshida et al. 2003; Hiltunen et al. 2014), thus providing us with an empirical test of this hypothesis.

We obviously did not impose any restriction on the range of colonization rates that could evolve. This may be unrealistic and other selective forces or constraints could of course interfere with the effects presented here. For instance, colonization rates may have upper bounds (see Supplementary Material for an example resulting from maximum reproductive investment). If the upper bound is low enough, this could constrain the possibility of at least some trophic levels to persist through the evolution of greater colonization rates. Large colonization rates are especially selected for at higher trophic levels, so the latter would need to have less stringent bounds.

One direct way to test our predictions is to use an experimental approach, with suitable biological systems such as ciliates or other microorganisms in which both dispersal and competition have been thoroughly studied (Fox & Morin, 2001; Pennekamp et al. 2014) and for which the definition of trophic levels, patches and dispersal are simple to establish. More generally, we might use observational data from the field to evaluate whether some of the qualitative patterns / trends we have identified here actually occurs – these patterns could indeed be diagnostic of whether the type of eco-evolutionary dynamics modelled are at play or not. Of course, natural communities and food chains experience processes and contingencies left out in models (*e.g.* autocorrelation and/or clustering of perturbations, species occurring at more than one trophic level). This would often complicate the task, but datasets such as the one used in Wang et al. (2021) exist and are extremely useful in this context, especially if they extend to three or more trophic levels. The patterns reported in Fig. 3 and 5 appear especially valuable for empirical assessments. Indeed, we found that among systems differing in their level of habitat destruction, either spontaneously or upon experimental manipulation, the maximum food-chain length declines in a generically convex manner in the absence of eco-evolutionary dynamics, whereas it could have a concave decline with such dynamics (Fig. 3c). Such a concave shape is never observed without evolutionary dynamics, and is not observed when manipulating the extinction rate. The existence of a concave relationship would therefore be a strong indication that eco-evolutionary dynamics are at play.

Alternatively, the correlation between total occupancy (and thus the fraction of unoccupied patches) and maximum food-chain length offers the same prediction (Fig. 5): it could assume a concave shape only with eco-evolutionary dynamics, not in their absence, and only if habitat destruction is the underlying varying parameter. A concave shape is observed upon varying the extinction rate *e* or the perturbation rate *μ* (Supplementary Fig. 2). It thus seems that datasets or experiments documenting the impact of habitat destruction on maximum food-chain length and overall occupancy offer great promises to evaluate the importance of eco-evolutionary dynamics in shaping food webs.

Our results were derived in a simple spatially homogeneous and mean-field patch-occupancy model. Interestingly, such a simple model faithfully reproduces the results presented in Wang et al. (2021), who used stochastic simulations of a spatially realistic model. This gives additional credence in the predictions we derived here. Our results could be extended in many different ways, possibly including more complex food-web topologies (“how would omnivory affect model predictions?”), introducing several species per trophic level (leading to the study of apparent predation at the scale of the metacommunity) or even using trophic level-specific definitions of a patch (*e.g.* McCann et al. 2005), but they reveal some fundamental ways in which the evolution of dispersal and spatial food-webs dynamics are interconnected. In any case, they suggest a lot is to be gained from bringing more eco-evolutionary dynamics into the realm of food-web ecology.

## Supporting information

Supplementary Information (mathematical appendix)

## Author contributions

The initial idea of this work originated from FM during his PhD thesis with PJ and PD. VC and FM designed and conducted the research. VC and FM wrote the initial manuscript, all co-authors contributed to manuscript preparation.

## Acknowledgments

VC acknowledges funding from INRAE, PJ, PD and FM from the CNRS. All authors acknowledge funding from ANR (project AFFAIRS 12-ADAP-005 to PD). VC and FM are thankful to CESAB, Montpellier, for hosting a work session. FM also thanks D. Gravel and various people at ISA and EDYSAN for feedback. VC dedicates the derivation of the function from a spatially structured population model to the dearly missed Isabelle Oliveri, who hinted at it in a split sentence in 2005, as he was doing his PhD under her supervision.

