## Supplementary Information (mathematical appendix) for "Coevolution of species colonization rates controls food-chain length in spatially structured food webs"

| Variable | Meaning |
|---|---|
| $p_i$ | Fraction of patches occupied by exactly *i* trophic levels. |
| $q_i$ | Fraction of patches occupied by at least *i* trophic levels; when focusing on a single trophic level, the index is dropped. |
| $h$ | When focusing on a single trophic level, denotes the fraction of patches occupied by its prey. |
| $s$ | When focusing on a single trophic level, denotes the fraction of patches occupied by its predator. |

| Parameter | Meaning | Typical value |
|---|---|---|
| $c_i$ | Colonization rate of trophic level *i*, which can evolve as the result of competition within the trophic level. | |
| $e$ | Rate at which a single trophic level goes extinct. | |
| $\mu$ | Perturbation rate, i.e. rate at which all trophic levels are wiped out of a patch. | |
| $q_0$ | Habitat availability for the first trophic level, i.e. fraction of patches that can actually be colonized by the first trophic level (the rest is considered non habitable). | 1 |
| $e^{TD}$ | Top-down control of extinction rates: the rate *e* is increased by this value for a trophic level in a patch in which its predator is also present. | |
| $\delta^{TD}$ | Relative top-down control of colonization rates: the colonization rate of a trophic level is multiplied by $1+\delta^{TD}$ in patches where its predator is also present. | |
| $\eta_{xy}$ | Probability that a colonist from species *x* replaces species *y* in a patch that the latter species occupies. This probability is linked to colonization rates of both species through the competition-colonization trade-off. | |

| Useful quantity | Meaning / usefulness | Formula |
|---|---|---|
| $\langle c_i \rangle$ | Average colonization rate of trophic level *i* over all the types of patches that it inhabits. | $\langle c_i \rangle = c_i \left(1+\frac{q_{i+1}}{q_i}\delta^{TD}\right)$ |
| $\langle e_i \rangle$ | Average trophic-level extinction rate of trophic level *i* over all the types of patches that it inhabits. | $\langle e_i \rangle = ie + \left(i-1+\frac{q_{i+1}}{q_i}\right)e^{TD}$ |
| $u_i$ | Helps write equilibrium occupancies. | $u_i = \sum_{j=1}^{i} \frac{1}{c_j}$ |
| $v_i$ | Helps write equilibrium occupancies. | $v_i = \sum_{j=1}^{i} \frac{j}{c_j}$ |
| $a_i$ | Coefficient associated to the first-order effect of $e^{TD}$ on occupancy of trophic level *i*. | eq. (1.10a) |
| $b_i$ | Coefficient associated to the first-order effect of $\delta^{TD}$ on occupancy of trophic level *i*. | eq. (1.10b) |
| $\epsilon_i$ | Quasi-exhaustive extinction rate of trophic level *i*: it includes all extinction rate terms except the one accounting for top-down control of the extinction rate of trophic level *i*. | $\epsilon_i = \mu + ie + (i-1)e^{TD}$ |
| $\alpha_i$ | Fraction of habitats occupied by trophic level *i* which are also occupied by its predator. | $\alpha_i = s_i / q_i = q_{i+1} / q_i$ |
| $r_y$ | Invasion fitness of species *y* competing against resident species *x*. | eq. (2.6) |
| $\beta_i$ | Multiplication of colonization rate of trophic level *i* due to top-down effects. | $\beta_i = 1 + \delta^{TD}\alpha_i$ |
| $\zeta_i$ | Exhaustive collection of extinction rates affecting trophic level *i*. | $\zeta_i = \epsilon_i + e^{TD}\alpha_i$ |
| $\phi$ | Intercept of the function $\eta$ when its two arguments are equal; measures the probability of neutral replacement in a patch. | $\phi = \eta(c,c)$ |
| $\psi(c)$ | Slope of the function $\eta$ with respect to the colonization rate of the species trying to take over a patch; defines the harshness of competition between species in the focal trophic level. | $\psi(c) = \frac{\partial \eta}{\partial x_1}(c,c)$ |
| $\omega(c)$ | Selection gradient of the invasion fitness. | eq. (2.11) |
| $\xi(c)$ | Second derivative of the function $\eta$ with respect to the colonization rate of the species trying to take over a patch; defines the "optimality" of singular colonization rates. | $\xi(c) = \frac{\partial^2 \eta}{\partial x_1^2}(c,c)$ |
| $\theta$, $\Theta$ | Generic names for single-variable functions describing the shape of function $\eta$. | |
| $\psi_0$, $\xi_0$ | Parameters used to describe functions $\psi$ and $\xi$ under particular assumptions. | |

### 1. Patch occupancy dynamics

#### *1.1. General spatialised food chain model*

As in Calcagno et al. (2011), we consider a simplified version of the spatialised food chain model described by Holt (1997, 2002), based on the metapopulation model of Levins & Culver (1971). Species/trophic level 1 refers to the species at the basis of the trophic chain, species/trophic level 2 is the predator of trophic level 1, and so forth. A species other than that at trophic level 1 cannot persist in a patch if there is no species from the previous trophic level. Therefore, when species *i* is present in a patch, all inferior species (1 to *i*-1) must also be present in the same patch.

Let $p_0$ be the proportion of empty patches in the metapopulation, and let $p_i$ be the proportion of patches occupied by species 1 to *i* and not by species *i*+1. We note $q_i$ the regional frequency of species *i* ($q_i = \sum_{j \geq i} p_j$), i.e. the proportion of patches occupied by *i* or more trophic levels. The rate of colonization by species i of patches already occupied by the first *i* – 1 species is proportional to its colonization ability ($c_i$) and to $q_i$ (each occupied patch contributes to the pool of potential colonizers through migrant production).

To give a simple illustration, the dynamics of a two-level system, in the absence of top-down control of colonization or extinction rates, are described by the following equations (Supp. Fig. 1):

$$\frac{dp_0}{dt} = (\mu + e_1) q_1 - c_1 p_0 q_1 \qquad (1.1a)$$

$$\frac{dp_1}{dt} = c_1 p_0 q_1 + e_2 q_2 - (\mu + e_1) p_1 - c_2 p_1 q_2 \qquad (1.1b)$$

$$\frac{dp_2}{dt} = c_2 p_1 q_2 - (\mu + e_1 + e_2) p_2 \qquad (1.1c)$$

where $c_1$ is the colonization rate of trophic level 1, $c_2$ that of trophic level 2, $\mu$ the rate at which perturbations happen which wipe all species from a patch at once, $e_1$ and $e_2$ the species-specific extinction rates of trophic levels 1 and 2 respectively.

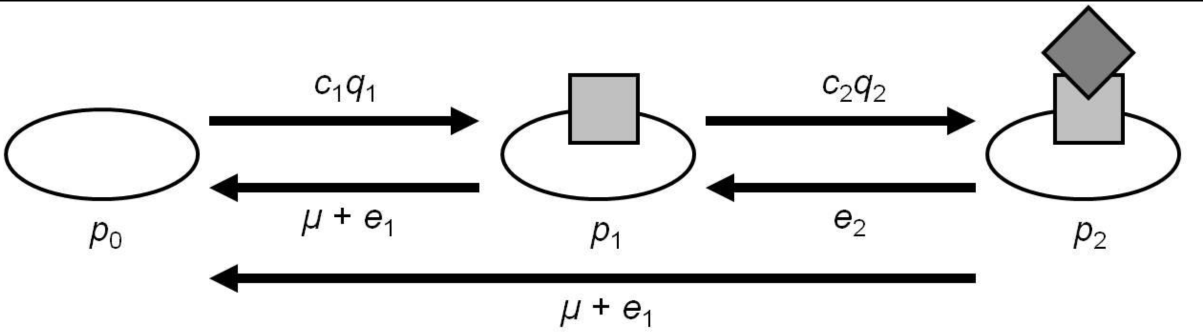

**Supp. Fig. 1** – Dynamics of the two-levels chain represented by system (1.1). Grey squares and diamonds correspond to the occurrence of the first and second-level species, respectively. Arrows represent flows of site frequency from one state to another and the formulae above them give the rates corresponding to these flows.

With top-down control of colonization rates, the term $c_1 q_1$ in equations (1.1a-b) would in fact become $c_1 p_1 + c_1^{TD} p_2$ where $c_1^{TD}$ would be the colonization rate of trophic level 1 from patches in which trophic level 2 is also present. In the same way, top-down control of extinction rates would turn the term $e_1 p_2$ in equation (1c) into $e_1^{TD} p_2$ and the term $e_1 q_1$ in equation (1a) into $e_1 p_1 + e_1^{TD} p_2$, with $e_1^{TD}$ the level-specific extinction rate of trophic level 1 in patches where trophic level 2 is also present.

It is generally more compact to write dynamics in terms of $q_i$, as in Calcagno et al. (2011):

$$\frac{dq_i}{dt} = \langle c_i \rangle q_i \left( q_{i-1} - q_i \right) - \left( \mu + \langle e_i \rangle \right) q_i \qquad \text{with } i > 0 \tag{1.2}$$

where $\langle c_i \rangle$ is used for the average colonization rate of trophic level *i* over all the types of patches that it inhabits (thus implicitly accounting for top-down control of colonization rate) and $\langle e_i \rangle$ denotes the average trophic-level extinction rate, also averaged over all the types of patches it inhabits.

#### *1.2. Model simplification*

To simplify parameterization, it is usually convenient to consider the case in which all trophic level-specific extinction rates are equal to *e*, all extinction rates in the presence of top-down effect are $e + e^{TD}$ and all top-down control of colonization rates can be summarized by coefficient $\delta^{TD}$, such that $c_i^{TD} = c_i \left( 1 + \delta^{TD} \right)$. Under these assumptions, the following relationships hold:

$$\langle e_i \rangle = ie + \left( i - 1 + \frac{q_{i+1}}{q_i} \right) e^{TD} \tag{1.3}$$

$$\langle c_i \rangle = c_i \left( 1 + \frac{q_{i+1}}{q_i} \delta^{TD} \right) \tag{1.4}$$

#### *1.3. Equilibrium occupancy*

The assumptions described by equations (1.3) and (1.4) lead to a particularly simple recursion system to solve to obtain the values of $q_i$ at equilibrium (i.e. setting the right-hand side of equation [1.2] equal to 0):

$$q_i = q_{i-1} - \frac{\mu + ie + \left( i - 1 + \frac{q_{i+1}}{q_i} \right) e^{TD}}{c_i \left( 1 + \frac{q_{i+1}}{q_i} \delta^{TD} \right)} \tag{1.5}$$

In the absence of top-down effects, the recursion can be explicitly computed:

$$q_i^* = q_0 - \sum_{j=1}^{i} \frac{\mu + je}{c_j} = q_0 - \mu u_i - e v_i \tag{1.6}$$

where quantities $u_i$ and $v_i$ are defined as:

$$u_i = \sum_{j=1}^{i} \frac{1}{c_j} \tag{1.7a}$$

$$v_i = \sum_{j=1}^{i} \frac{j}{c_j} \tag{1.7b}$$

In the simplest case, one can assume $q_0 = 1$, i.e. all patches are available for colonization by the first trophic level. However, keeping in line with Nee & May (1992), we will consider $q_0$ in all generality.

Assuming that top-down control of colonization and extinction rates is weak ( $e^{TD} \ll e$ and $\delta^{TD} \ll 1$), we can approximate the result of recursion (1.5) as a first-order modification of equation (1.6):

$$q_i^* \approx q_0 - \mu u_i - ev_i + a_i e^{TD} + b_i \delta^{TD} \quad (1.8)$$

with coefficients $a_i$ and $b_i$ obtained through the following recursions:

$$a_i = a_{i-1} - \frac{1}{c_i}\left[ i - 1 + \frac{q_0 - \mu u_{i+1} - ev_{i+1}}{q_0 - \mu u_i - ev_i} \right] \quad (1.9a)$$

$$b_i = b_{i-1} + \frac{\mu + ie}{c_i} \frac{q_0 - \mu u_i - ev_i}{q_0 - \mu u_i - ev_i} \quad (1.9b)$$

System (1.9) can be solved, setting $a_0 = 0$ and $b_0 = 0$ as initial conditions:

$$a_i = -\sum_{j=1}^{i} \frac{1}{c_j}\left[ j - 1 + \frac{q_0 - \mu u_{j+1} - ev_{j+1}}{q_0 - \mu u_j - ev_j} \right] = u_i - v_i - \sum_{j=1}^{i} \frac{1}{c_j}\left[ \frac{q_0 - \mu u_{j+1} - ev_{j+1}}{q_0 - \mu u_j - ev_j} \right] \quad (1.10a)$$

$$b_i = \sum_{j=1}^{i} \left( \frac{\mu + je}{c_j} \right) \left( \frac{q_0 - \mu u_{j+1} - ev_{j+1}}{q_0 - \mu u_j - ev_j} \right) \quad (1.10b)$$

A more precise way of solving for the equilibrium occupancy when top-down control exists is to rewrite equation (1.2) in vector and matrix form:

$$\frac{d\vec{q}}{dt} = \begin{bmatrix} 0 & 0 & \cdots & \cdots & 0 \\ \langle c_1 \rangle q_1 & -\left(\mu + \langle e_1 \rangle + \langle c_1 \rangle q_1\right) & \ddots & \ddots & \vdots \\ 0 & \langle c_2 \rangle q_2 & \ddots & \ddots & \vdots \\ \vdots & \ddots & \ddots & \ddots & 0 \\ 0 & \cdots & 0 & \langle c_S \rangle q_S & -\left(\mu + \langle e_S \rangle + \langle c_S \rangle q_S\right) \end{bmatrix} . \vec{q} \quad (1.11)$$

A reliable algorithm is, starting from an informed guess on the value of vector $\vec{q}$ (with first element $q_0$ fixed as a model parameter), to look for the eigenvector associated with eigenvalue 0 for the matrix in equation (1.11), and then to renormalize the eigenvector so that its first element is exactly $q_0$. After some iterations, the algorithm converges towards the equilibrium occupancy vector, provided that the top-down effects are not too strong. Indeed, the iterations between successive vectors $\vec{q}^{\,t}$ are given by the following recursion:

$$q_n^{t+1} = q_0 \frac{\prod_{i=1}^{n} \langle c_i^t \rangle q_i^t}{\prod_{i=1}^{n} \left[ \mu + \langle e_i^t \rangle + \langle c_i^t \rangle q_i^t \right]} \quad (1.12)$$

where $\langle e_i^t \rangle$ and $\langle c_i^t \rangle$ are the effective extinction and colonization rates of trophic level *i* at iteration *t* (corresponding to equations [1.3] and [1.4]). Observing that the equilibrium vector $\vec{q}^{\,*}$ also obeys a similar equation:

$$q_n^* = q_0 \frac{\prod_{i=1}^{n} \langle c_i^* \rangle q_i^*}{\prod_{i=1}^{n} \left[ \mu + \langle e_i^* \rangle + \langle c_i^* \rangle q_i^* \right]} \tag{1.13}$$

in which all quantities with stars correspond to their equilibrium values, the recursion amongst the deviations from the equilibrium can be computed. For instance, the recursion of the deviation $\Delta q_1$ for $q_1$ can be written:

$$\Delta q_1^{t+1} = \frac{q_0 - q_1^*}{q_0} \Delta q_1^t + \frac{c_1 \left(q_0 - q_1^*\right) - \mu - e}{\langle c_1^* \rangle q_0} \left( \Delta q_1^t - \frac{q_1^*}{q_2^*} \Delta q_2^t \right) \tag{1.14}$$

The second group of terms on the right-hand side of (1.14) can be easily controlled if $\delta^{TD} \ll 1$ and $e^{TD} \ll e$. In the same vein, the other deviations display similar recursions, which converge towards zero when top-down effects are dominated by the other parameters.

### 2. Adaptive dynamics

#### *2.1. Generic occupancy dynamics*

Trying to simplify the dynamics of system (1.2), one might consider that for any trophic level $n$, the dynamics of its occupancy $q_n$ depends on the proportion of patches it does not yet occupy but can colonize (fraction $h_n = q_{n-1}$) and on the proportion of patches that are already colonized by the focal trophic level and its immediate predator (fraction $s_n = q_{n+1}$). The equivalent of equation (1.2) becomes:

$$\frac{dq}{dt} = cq\left(1 + \frac{s}{q}\delta^{TD}\right)(h - q) - \left[\mu + ne + \left(n - 1 + \frac{s}{q}\right)e^{TD}\right]q \tag{2.1}$$

where all *n* indices have been omitted for the sake of clarity.

Using compound parameter $\epsilon_n = \mu + ne + (n-1)e^{TD}$, equation (2.1) can be further simplified as:

$$\frac{dq}{dt} = c\left(q + \delta^{TD} s\right)(h - q) - \epsilon q - e^{TD} s \tag{2.2}$$

Solving for the equilibrium of equation (2.2) yields an equation equivalent to equation (1.5):

$$q = h - \frac{\epsilon + e^{TD} s / q}{c\left(1 + \delta^{TD} s / q\right)} \tag{2.3}$$

Finally, the value of $\alpha_n = s_n / q_n$ used in equation (2.3), and more generally whenever it is multiplied by top-down effect parameters, can be obtained using equation (1.6), i.e. only keeping the $0^{\text{th}}$ order approximation of the fraction with respect to top-down effect parameters:

$$\alpha_n = \frac{q_{n+1}}{q_n} \approx \frac{q_0 - \mu u_{n+1} - e v_{n+1}}{q_0 - \mu u_n - e v_n} = 1 - \frac{1}{c_{n+1}} \frac{\mu + (n+1)e}{q_0 - \mu u_n - e v_n} \tag{2.4}$$

#### *2.2. The competition-colonization model*

We now consider the dynamics of two species occupying the same trophic level, using the formalism of Calcagno et al. (2006) to generalize the very strict trade-off between colonization rate and competitiveness implemented by Tilman (1994). We consider two species *x* and *y* such that their respective occupancies follow:

$$\frac{dq_x}{dt} = c_x\left(q_x + \delta^{TD} s_x\right)\left(h_n - q_x - q_y\right) - \epsilon_n q_x - e^{TD} s_x \\ + \eta_{xy} c_x\left(q_x + \delta^{TD} s_x\right) q_y - \eta_{yx} c_y\left(q_y + \delta^{TD} s_y\right) q_x \quad (2.5a)$$

$$\frac{dq_y}{dt} = c_y\left(q_y + \delta^{TD} s_y\right)\left(h_n - q_x - q_y\right) - \epsilon_n q_y - e^{TD} s_y \\ - \eta_{xy} c_x\left(q_x + \delta^{TD} s_x\right) q_y + \eta_{yx} c_y\left(q_y + \delta^{TD} s_y\right) q_x \quad (2.5a)$$

where $\eta_{xy}$ (resp. $\eta_{yx}$) is the probability that a colonist from species *x* (resp. *y*) takes over a patch already occupied by species *y* (resp. *x*). In the absence of top-down effect, both species experience the same extinction rate $\epsilon_n$.

In the following, we assume that all trophic levels are at ecological equilibrium (i.e. such that the right-hand side of equation [1.2] equals 0 for all trophic levels), but within a particular trophic level, we are interested in the dynamics of two different species competing for dominance within this trophic level, and thus do not assume equilibrium within the trophic level. If the immediately upper trophic level does not have more than one species, developing the ODEs for $s_x$ and $s_y$ and assuming quasi-equilibrium yields $s_x = \alpha_n q_x$ and $s_y = \alpha_n q_y$. Equation (2.5a) divided by $q_y$ and simplified by the proportionality between $s_y$ and $q_y$ leads to the following invasion fitness $r_y$:

$$r_y = \frac{1}{q_y}\frac{dq_y}{dt}\bigg|_{\substack{q_y=0 \\ q_x=q}} = c_y \beta_n\left(h_n - q\right) - \zeta_n + \beta_n\left(\eta_{yx} c_y - \eta_{xy} c_x\right) q \quad (2.6)$$

with $\beta_n = 1 + \delta^{TD}\alpha_n$ and $\zeta_n = \epsilon_n + e^{TD}\alpha_n$.

In order to explicit the relationship between competitiveness and colonization rates, we will further assume that the probabilities $\eta_{xy}$ and $\eta_{yx}$ are obtained through a function $\eta$ of two variables, such that:

$$\eta_{xy} = \eta\left(c_x, c_y\right) \quad (2.7a)$$

$$\eta_{yx} = \eta\left(c_y, c_x\right) \quad (2.7b)$$

This function can be trophic level-specific, if needed. We will generically write $\eta(x_1, x_2)$ to explicit the first and second variables of function $\eta$.

#### *2.3. Selection gradient and singular strategies*

The first derivative of the invasion fitness $r_y$ with respect to the colonization rate $c_y$ of the rare type is given by:

$$\frac{\partial r_y}{\partial c_y} = \beta_n\left[h_n - q + \left(\eta\left(c_y, c_x\right) + \frac{\partial \eta}{\partial x_1} c_y - \frac{\partial \eta}{\partial x_2} c_x\right) q\right] \quad (2.8)$$

Equation (2.7) can be simplified by plugging in equation (2.3), i.e. $q = h_n - \zeta_n / c_x \beta_n$:

$$\frac{\partial r_y}{\partial c_y} = \frac{1}{c_x}\left[\zeta_n + \left(\eta\left(c_y, c_x\right) + \frac{\partial \eta}{\partial x_1} c_y - \frac{\partial \eta}{\partial x_2} c_x\right)\left(h_n c_x \beta_n - \zeta_n\right)\right] \tag{2.9}$$

To go further, we need to make assumptions on function $\eta$. A first, reasonable assumption is to assume that neutral competitiveness does not depend on the colonization rate, i.e. that $\eta\left(c,c\right)$ is independent of *c*. Taking the first derivative of this relation with respect to *c*, we obtain:

$$\frac{\partial \eta}{\partial x_1}\left(c,c\right) + \frac{\partial \eta}{\partial x_2}\left(c,c\right) = 0 \tag{2.10}$$

If we define $\phi = \eta\left(c,c\right)$ and $\psi\left(c\right) = \frac{\partial \eta}{\partial x_1}\left(c,c\right)$, the selection gradient $\omega(c)$ of $c$ can be obtained by plugging assumptions on $\eta$ into equation (2.9):

$$\omega\left(c\right) = \left.\frac{\partial r_y}{\partial c_y}\right|_{\substack{c_y \to c_x \\ c_x = c}} = \frac{1}{c}\left[\zeta_n + \left(\phi + 2\psi\left(c\right)c\right)\left(h_n c \beta_n - \zeta_n\right)\right] \tag{2.11}$$

Values of $c$ at which the selection gradient vanishes are singular strategies in the adaptive dynamics parlance (Geritz et al., 1998), defined by the implicit equation:

$$\zeta_n + \left(\phi + 2\psi\left(c\right)c\right)\left(h_n c \beta_n - \zeta_n\right) = 0 \tag{2.12}$$

Because we assume a trade-off between colonization rate and competitiveness, $\psi(c) < 0$ for all values of $c$. The equation (2.12) can be re-arranged to illustrate the fact that, at singular strategies, the marginal benefits of being a better colonizer (given by the proportion of patches still devoid of the focal trophic level, $\zeta_n / c\beta_n$ and the opportunity of neutral replacements $\phi q$) exactly balance the marginal cost of decreased competitive ability multiplied by the proportion of patches already occupied by the focal trophic level:

$$\frac{\zeta_n}{c\beta_n} + \phi q = -2\psi\left(c\right)cq \tag{2.13}$$

We can also note in passing that if $\psi(c)$ is a polynomial of $c$ of degree $d$, the singular strategy condition (2.12) admits at most $d+1$ solutions.

#### *2.4. Convergence stability*

A singular strategy is said to be convergence stable (CS) if and only if directional selection leads to the singular strategy (Geritz et al., 1998). This means that the derivative of the selection gradient is negative at the singular strategy, i.e.

$$\frac{d\omega}{dc} = \frac{1}{c^2}\left\{2c^2\left[\left(h_n c\beta_n - \zeta_n\right)\psi'(c) + h_n\beta_n\psi(c)\right] - \zeta_n\left(1-\phi\right)\right\} < 0 \tag{2.14}$$

with

$$\psi'(c) = \frac{\partial^2 \eta}{\partial x_1^2}\left(c,c\right) + \frac{\partial^2 \eta}{\partial x_1 \partial x_2}\left(c,c\right) \tag{2.15}$$

We immediately notice that, since $\psi(c) < 0$, $0 < \phi < 1$ and $h_n c\beta_n - \zeta_n = qc\beta_n > 0$, a singular strategy $c$ is CS if and only if

$$\psi'(c) < \frac{1}{h_n\beta_n c - \zeta_n}\left(\frac{\zeta_n(1-\phi)}{2c^2} - h_n\beta_n\psi(c)\right) \tag{2.16}$$

The parameter $\zeta_n$ can be eliminated from condition (2.16) by using the singular strategy condition (2.12):

$$\psi'(c) < \frac{2}{c^2}\left[\psi^2c^2 - (1-\phi)\psi c - \frac{\phi(1-\phi)}{4}\right] \tag{2.17}$$

The polynomial function of $\psi c$ in the right-hand side of condition (2.17) attains a minimum at $c\psi(c) = (1-\phi)/2$ and the value of the right-hand side at this minimum is $-(1-\phi)/2c^2$. However, $c\psi(c) = (1-\phi)/2$ is not attainable given that $\psi(c) < 0$. The two roots of the polynomial are of opposite signs, so the minimum attainable value of the right-hand side of condition (2.17) is obtained when $\psi(c) = 0$ and is equal to $-\phi(1-\phi)/2c^2$.

We note in passing that $\psi'(c)$ can be rewritten (developing equation [2.15] using equation [2.10]):

$$\begin{aligned}\psi'(c) &= \frac{\partial^2\eta}{\partial x_1^2}(c,c) + \frac{\partial^2\eta}{\partial x_1\partial x_2}(c,c) \\ &= -\frac{\partial^2\eta}{\partial x_2^2}(c,c) - \frac{\partial^2\eta}{\partial x_1\partial x_2}(c,c) \\ &= \frac{1}{2}\left[\frac{\partial^2\eta}{\partial x_1^2}(c,c) - \frac{\partial^2\eta}{\partial x_2^2}(c,c)\right]\end{aligned} \tag{2.18}$$

For the sake of clarity, we will note $\xi(c) = \frac{\partial^2\eta}{\partial x_1^2}(c,c)$. The three second derivatives can be rewritten using $\psi'(c)$ and $\xi(c)$:

$$\frac{\partial^2\eta}{\partial x_1^2}(c,c) = \xi(c) \tag{2.19a}$$

$$\frac{\partial^2\eta}{\partial x_1\partial x_2}(c,c) = \psi'(c) - \xi(c) \tag{2.19b}$$

$$\frac{\partial^2\eta}{\partial x_2^2}(c,c) = \xi(c) - 2\psi'(c) \tag{2.19c}$$

***2.5. Evolutionary stability***

A singular strategy is called an evolutionarily stable strategy (ESS) if and only if it cannot be invaded by similar strategies. In practice, this amounts to checking whether the second derivative of the invasion fitness with respect to the colonization rate of the rare type is negative (Hofbauer and Sigmund, 1990, Geritz et al., 1998), i.e. that

$$\left.\frac{\partial^2 r_y}{\partial c_y{}^2}\right|_{c_x=c_y=c} = \frac{1}{c}\left(2\psi(c)+2\xi(x)c-\psi'(c)c\right)\left(h_n c\beta_n - \zeta_n\right) < 0 \tag{2.20}$$

Given that $h_n c\beta_n - \zeta_n > 0$, the ESS condition can be rewritten as:

$$2\psi(c)+2\xi(x)c-\psi'(c)c < 0 \tag{2.21}$$

***Conclusions of the generic adaptive dynamics***

For the considered trophic level $n$, the condition to find singular strategies $c$ is:

$$\zeta_n + \left(\phi + 2\psi\left(c\right)c\right)\left(h_n c\beta_n - \zeta_n\right) = 0$$

A singular strategy is CS if and only if:

$$\psi'(c) < \frac{2}{c^2}\left[\psi(c)^2 c^2 - \left(1-\phi\right)\psi(c)c - \frac{\phi\left(1-\phi\right)}{4}\right]$$

A singular strategy is ESS if and only if:

$$2\psi(c)+2\xi(x)c-\psi'(c)c < 0$$

### 3. Applications

Here, we apply the adaptive dynamics conditions obtained above under particular cases of function $\eta$.

#### *3.1. Multiplicative $\eta$ function*

Here, we assume that

$$\eta\left(x_1, x_2\right) = \theta\left(x_1 / x_2\right) \tag{3.1}$$

for a certain, well-behaved continuous decreasing function $\theta$, i.e. the probability that species 1 takes over a patch where species 2 is present depends on the ratio $c_1 / c_2$. This implies that a species that has half the colonization rate of its competitor has a probability of taking over a patch which does not depend on the colonization rate of the "better colonizer" – as long as the ratio is ½, the probability of taking over stays the same. Defining $\Theta = \theta \circ \exp$ as a composed function based on $\theta$ (with naturally $\theta = \Theta \circ \log$), we easily see that the set of all $\eta$ functions defined by equation (3.1) is also the same as the set of $\eta$ function defined as functions of the difference of the log of its variables:

$$\eta\left(x_1, x_2\right) = \Theta\left(\log(x_1) - \log(x_2)\right) \tag{3.2}$$

These definitions of $\eta$ entail that

$$\psi(c) = \frac{\partial \eta}{\partial x_1}(c,c) = \frac{1}{c}\frac{d\Theta}{dx}(0) \tag{3.3a}$$

i.e. $\psi(c)$ can be written as a hyperbolic function , and

$$\xi(c) = \frac{\partial^2 \eta}{\partial x_1^2}(c,c) = \frac{1}{c^2}\left[\frac{d^2\Theta}{dx^2}(0) - \frac{d\Theta}{dx}(0)\right] \tag{3.3b}$$

We will here use the following forms:

$$\psi(c) = \frac{\psi_0}{c} \tag{3.4a}$$

$$\xi(c) = \frac{\xi_0}{c^2} \tag{3.4b}$$

Because $\Theta$ is decreasing, $\psi_0 < 0$. However, depending on the relative convexity of function $\Theta$, $\xi_0$ can be positive or negative.

For this particular case, the singular strategy equation becomes:

$$\zeta_n + \left(\phi + 2\psi_0\right)\left(h_n c \beta_n - \zeta_n\right) = 0 \tag{3.5}$$

which yields a unique solution:

$$c = \frac{\zeta_n}{h_n \beta_n}\left(1 - \frac{1}{\phi + 2\psi_0}\right) \tag{3.6}$$

The value of $c$ given by equation (3.6) is non-negative when

$$\phi > 1 - 2\psi_0 \tag{3.7}$$

or

$$\phi < -2\psi_0 \tag{3.8}$$

However, we see that when $\phi + 2\psi_0 > 0$, all the components of equation (3.5) are positive (if we assume $q > 0$), which should lead to selection for infinite colonization rates. So we need $\phi < -2\psi_0$ for this case to be biologically relevant.

Provided parameters $\phi$ and $\psi_0$ are shared by all trophic levels, an explicit solving of recursion (1.5) in the absence of top-down effects, together with equation (3.6) yields:

$$q_i^* = \frac{q_{i-1}}{1 - \phi - 2\psi_0} = \frac{q_0}{\left(1 - \phi - 2\psi_0\right)^i} \tag{3.9}$$

with ESS colonization rates set at:

$$c_i^{ESS} = \left(\mu + ie\right)\frac{\left(1 - \phi - 2\psi_0\right)^i}{-\left(\phi + 2\psi_0\right)} \tag{3.10}$$

Equation (3.9) indicates that, with this form of the trade-off function, occupancies are independent of perturbation and extinction rates, and are only determined by the availability of patches and the parameters of the trade-off curve. Equation (3.10) also shows that the product $c_i^{ESS} q_i^*$ increases linearly with $i$.

In the presence of small but non-zero top-down effects, the equilibrium occupancies become:

$$q_i^* \approx \frac{q_0}{\left(1 - \phi - 2\psi_0\right)^i} + a_i e^{TD} + b_i \delta^{TD} \tag{3.11}$$

while the ESS colonization rates are approximated as:

$$c_i^{ESS} \approx \left(\mu + ie\right)\frac{\left(1 - \phi - 2\psi_0\right)^i}{-\left(\phi + 2\psi_0\right)} + f_i e^{TD} + g_i \delta^{TD} \tag{3.12}$$

with $a_i$, $b_i$, $f_i$ and $g_i$ given by:

$$a_i = -\left(\frac{\phi + 2\psi_0}{1-\phi-2\psi_0}\right)u_i - v_i \tag{3.13a}$$

$$b_i = \frac{\mu}{1-\phi-2\psi_0}u_i + \frac{e}{1-\phi-2\psi_0}v_j \tag{3.13b}$$

$$f_i = \frac{(1-\phi-2\psi_0)^i}{-(\phi+2\psi_0)}\left(i-1+\frac{1}{1-\phi-2\psi_0}\right) \tag{3.13c}$$

$$g_i = (\mu + ie)\frac{(1-\phi-2\psi_0)^{i-1}}{\phi+2\psi_0} \tag{3.13d}$$

$$\begin{aligned} u_i &= -(\phi+2\psi_0)\sum_{j=1}^{i}\frac{1}{(\mu+je)(1-\phi-2\psi_0)^j} \\ &= -\frac{(\phi+2\psi_0)}{e(1-\phi-2\psi_0)}\left\{\begin{array}{l}\Phi\left[\frac{1}{1-\phi-2\psi_0},1,1+\frac{\mu}{e}\right] \\ -\frac{1}{(1-\phi-2\psi_0)^i}\Phi\left[\frac{1}{1-\phi-2\psi_0},1,1+i+\frac{\mu}{e}\right]\end{array}\right\}\end{aligned} \tag{3.13e}$$

$$\begin{aligned} v_i &= -(\phi+2\psi_0)\sum_{j=1}^{i}\frac{j}{(\mu+je)(1-\phi-2\psi_0)^j} \\ &= \frac{1}{e^2}\left\{\begin{array}{l} e\left[1-\frac{1}{(1-\phi-2\psi_0)^i}\right] \\ +\mu\left(\frac{\phi+2\psi_0}{1-\phi-2\psi_0}\right)\left(\begin{array}{l}\Phi\left[\frac{1}{1-\phi-2\psi_0},1,1+\frac{\mu}{e}\right] \\ -\frac{1}{(1-\phi-2\psi_0)^i}\Phi\left[\frac{1}{1-\phi-2\psi_0},1,1+i+\frac{\mu}{e}\right]\end{array}\right)\end{array}\right\}\end{aligned} \tag{3.13f}$$

where $\Phi$ is the Lerch transcendent $\Phi[z,s,a] = \sum_{j=0}^{\infty}\frac{z^j}{(j+a)^s}$.

The singular strategy is CS if and only if:

$$-\psi_0 < 2\left[\psi_0^2 - (1-\phi)\psi_0 - \frac{\phi(1-\phi)}{4}\right] \tag{3.14}$$

which is always true when $\phi < -2\psi_0$, a condition required to have a non-negative singular strategy (equations [3.6-3.8]).

The singular strategy is ESS if and only if:

$$\xi_0 < -\frac{3\psi_0}{2} \tag{3.15}$$

#### *3.2. Additive $\eta$ function*

Here, we assume that

$$\eta(x_1, x_2) = \theta(x_1 - x_2) \tag{3.16}$$

for a certain, well-behaved continuous decreasing function $\theta$.
This definition of $\eta$ entails that

$$\psi(c) = \frac{\partial \eta}{\partial x_1}(c,c) = \frac{d\theta}{dx}(0) \tag{3.17a}$$

and

$$\xi(c) = \frac{\partial^2 \eta}{\partial x_1^2}(c,c) = \frac{d^2\theta}{dx^2}(0) \tag{3.17b}$$

i.e. both $\psi$ and $\xi$ are independent of $c$. We will here use the following forms:

$$\psi(c) = \psi_0 \tag{3.18a}$$

$$\xi(c) = \xi_0 \tag{3.18b}$$

For this particular case, the singular strategy equation becomes:

$$\zeta_n + (\phi + 2\psi_0 c)(h_n c\beta_n - \zeta_n) = 0 \tag{3.19}$$

which formally admits two solutions:

$$c_{\pm}^{ESS} = \frac{2\zeta_n\psi_0 - h_n\beta_n\phi \pm \sqrt{(h_n\beta_n\phi - 2\zeta_n\psi_0)^2 - 8h_n\beta_n\zeta_n(1-\phi)}}{4h_n\beta_n\psi_0} \tag{3.20}$$

However, only one of these solutions is positive (they are of opposite signs since $\psi_0 < 0$).

$$c_i^{ESS} = \frac{\zeta_i}{2q_{i-1}\beta_i} - \frac{\phi}{4\psi_0} - \frac{\sqrt{(q_{i-1}\beta_i\phi - 2\zeta_i\psi_0)^2 - 8q_{i-1}\beta_i\zeta_i(1-\phi)}}{4q_{i-1}\beta_i\psi_0} \tag{3.21}$$

where

$$\zeta_i = \mu + ie + \left(i - 1 + \frac{q_{i+1}}{q_i}\right)e^{TD} \tag{3.22}$$

and

$$\beta_i = 1 + \delta^{TD}\frac{q_{i+1}}{q_i} \tag{3.23}$$

To simplify the computation of equilibrium values for both $q_i$ and $c_i$, another strategy is to re-write the evolutionary equilibrium obtained in equation (3.19) as a function of the $q_i$:

$$q_{i-1} - q_i + (\phi + 2\psi_0 c_i)q_i = 0 \tag{3.24}$$

which leads to an expression linking $c_i$ to the $q_i$'s:

$$c_i^{ESS} = \frac{1}{2\psi_0}\left(1 - \phi - \frac{q_{i-1}}{q_i}\right) \tag{3.25}$$

Plugging this into equation (1.5) when top-down effects are present yields:

$$q_{i+1} = -q_i \frac{\left(1-\frac{q_i}{q_{i-1}}\right)\left(1-(1-\phi)\frac{q_i}{q_{i-1}}\right)+2\psi_0 \frac{q_i}{q_{i-1}} \frac{\mu+ie+(i-1)e^{TD}}{q_{i-1}}}{\left(1-\frac{q_i}{q_{i-1}}\right)\left(1-(1-\phi)\frac{q_i}{q_{i-1}}\right)\delta^{TD}+2\psi_0 \frac{e^{TD}}{q_{i-1}}} \tag{3.26}$$

When top-down control only affects the extinction rate, the recursion simplifies to:

$$q_{i+1} = -q_i q_{i-1} \frac{\left(1-\frac{q_i}{q_{i-1}}\right)\left(1-(1-\phi)\frac{q_i}{q_{i-1}}\right)+2\psi_0 \frac{q_i}{q_{i-1}} \frac{\mu+ie+(i-1)e^{TD}}{q_{i-1}}}{2\psi_0 e^{TD}} \tag{3.27}$$

When top-down control only affects the colonization rate, equation (3.26) becomes:

$$q_{i+1} = -\frac{q_i}{\delta^{TD}}\left[1+\frac{2\psi_0 \frac{q_i}{q_{i-1}} \frac{\mu+ie}{q_{i-1}}}{\left(1-\frac{q_i}{q_{i-1}}\right)\left(1-(1-\phi)\frac{q_i}{q_{i-1}}\right)}\right] \tag{3.28}$$

Plugging equation (3.25) into equation (1.5) in the absence of any top-down effect gives another, simpler recursion:

$$q_i = q_{i-1} \frac{2-\phi-2\psi_0 \frac{\mu+ie}{q_{i-1}} - \sqrt{\left(2-\phi-2\psi_0 \frac{\mu+ie}{q_{i-1}}\right)^2 - 4(1-\phi)}}{2(1-\phi)} \tag{3.29}$$

The singular strategy is CS if and only if:

$$0 < \frac{2}{c^2}\left[\psi_0^2 c^2 - (1-\phi)\psi_0 c - \frac{\phi(1-\phi)}{4}\right] \tag{3.30}$$

which is equivalent to:

$$c > \frac{1-\phi-\sqrt{1-\phi}}{2\psi_0} \tag{3.31}$$

However, given that $\psi_0 < 0$, equation (3.21) shows that $c > -\phi / 4\psi_0$, which proves that condition (3.31) is always true.

The singular strategy is ESS if and only if:

$$\xi_0 c < -\psi_0 \tag{3.32}$$

### 4. A mechanistic derivation of the additive $\eta$ function from a spatially structured population model

In the above, the existence of the patch-dynamic model and its function $\eta$ and the shape of the latter are postulated, not derived from a particular mechanism. As an alternative, let us consider a metapopulation consisting of a large number *N* of demes (local populations, or patches for that matter). Each patch, when occupied, can host a local population of size *n*. Assume that each individual produces dispersive propagules at some rate *d*, which are sent to another random deme, and local propagules at rate *b*, which stay to compete in the focal deme. Within a local deme, all propagules from all individuals compete for vacant sites according to a simple competitive lottery model.

Assume that dispersing propagules are more costly to produce than non dispersing ones, so that producing a disperser costs $\sigma$ as much energy as producing a non disperser. If the total energy investment in propagule production is $\Lambda$, then we have the linear trade-off

$$\Lambda = \sigma d + b \qquad (4.1)$$

At this stage, we must make some assumptions in order to align with the classic patch dynamics metapopulation scenario, that we call the Levins' limit. We first assume that reproduction is not limiting, i.e. $\Lambda$ is very large. This has two implications. First, there is no maximal value that *d* can take, i.e. *b* will never fall to zero for finite *d*. Second, as *b* is very large, this implies that the local population dynamics is very fast, and that local populations quickly go to their equilibrium size *n* and stay there, unless a perturbation causes the patch to go extinct (at the usual rate *e*).

Second, we assume that only a fraction $1-l$ of dispersers survive dispersal, while the others are lost in the process. In other words, *l* is the cost of dispersal. Under those conditions, the total rate of propagules arrival in a patch at a given time is proportional to occupancy, times the local population size *n*, times $d(1-l)$. We assume that *l* is sufficiently close to 1, so that few propagules manage to successfully land in patches. This guarantees that there will always be a fraction of unoccupied patches at any time, and also that immigration events do not contribute significantly to the local population dynamics (separation of time scales). If a propagule manages to land in an empty patch, since *b* is very large, it will always manage to establish a local population of size *n*, i.e. we can neglect the risk of initial stochastic extinction. Under these conditions, the colonization rate is

$$c = nd(1-l) \qquad (4.2).$$

Now if a disperser propagule of genotype *i* arrives in a patch already occupied by a local population of genotype *j*, the two genotypes will compete and one or the other will soon prevail (fixate), as local population dynamics is fast. If genotype *i* invests more in local reproduction, it has some chance to overtake the patch. Otherwise, it goes extinct. In a large population, the probability of fixation of an initially rare but slightly advantageous variant is well approximated by *s* (here given by $b_i - b_j$ for genotype *i*), where *s* is the selection coefficient, i.e. the advantage in local fitness of the favoured genotype (Haldane, 1927, Wright, 1931, Kimura, 1962). Therefore, the probability of fixation of genotype *i*, if it is advantaged locally, is

$$b_i - b_j = \sigma(d_j - d_i) = \frac{\sigma}{n(1-l)}(c_j - c_i) \qquad (4.3)$$

If genotypes vary in their colonization rates through their investment in dispersing versus resident propagules, we thus recover the additive $\eta$ function, with $\eta(c_i - c_j)$ given by

$$\eta_{i,j} = \eta\left(c_i - c_j\right) = \begin{cases} \dfrac{\sigma}{n\left(1-l\right)}\left(c_j - c_i\right) & \text{if } c_j > c_i \\ 0 & \text{otherwise} \end{cases} \tag{4.4}$$

In the case of two identical genotypes, $c_i = c_j$ and thus the immigrant genotype never overtakes the patch. This is consistent with the expression of the probability of fixation of rare neutral variant in a population of size *n*, which is $1/n$, and goes to zero as *n* gets large.

It follows that under this model, we obtain the special case

$$\phi = 0 \tag{4.5a}$$

$$\psi_0 = -\frac{\sigma}{n\left(1-l\right)} \tag{4.5b}$$

of the additive $\eta$ model, where $\psi_0$ is obtained by taking the left-derivative with respect to $c_i$. The larger the energetic cost of producing dispersing propagules, and the greater the cost of dispersal, the steeper the slope of the competitive trade-off.

A more general approximation exists for the probability of fixation in large populations, which allows us to consider the (small) possibility that a rare disadvantaged immigrant genotype can still overtake the patch by chance. We can indeed use the probability of fixation of an initially rare allele in a population of constant size *n*, which is well known in population genetics (Kimura, 1957). If the selection coefficient is *s* as defined above, then the probability of fixation of genotype *i* if it arrives in initial frequency $1/n$ in a population of size *n* is well approximated by (similarly to the allele fixation in the haploid case)

$$\eta(s) \approx \frac{1-e^{-2s}}{1-e^{-2ns}} \tag{4.6}$$

In the special case of two identical genotypes, $s = 0$, and the above expression is indeterminate. Using L'Hospital's rule, we find its value to be $1/n$, as expected. If we linearize to second order in *s* at $s = 0$, taking care of the indeterminacies, we obtain the approximation

$$\eta_{i,j} = \eta\left(c_i - c_j\right) \approx \frac{1}{n} + \frac{n-1}{n}\frac{\sigma}{n\left(1-l\right)}\left(c_j - c_i\right) \tag{4.7}$$

This leads to a result very similar to equations (4.5), with the following parameters:

$$\phi = 1/n \tag{4.8a}$$

$$\psi_0 = -\frac{n-1}{n}\frac{\sigma}{n\left(1-l\right)} \tag{4.8b}$$

This substantiates our interpretation of $\phi$ as a kin competition term, since in that case $\phi = 1/n$ is equal to the relatedness coefficient within a patch, i.e. the probability that an individual taken at random in the patch is identical by descent to a newly appeared individual in this patch. For instance, if we assume that the habitat is saturated by the species, with no unoccupied patches ($q_i = 1$), we see that $\phi$ still imposes some positive selection for dispersal, the more so when local population size ($n$) is small, mirroring a famous result on ES dispersal rate by Hamilton and May (1977).

The reader can refer to Jansen and Vitalis (2007) for similar derivations under slightly different assumptions.

### 5. Effect of the perturbation rate $\mu$

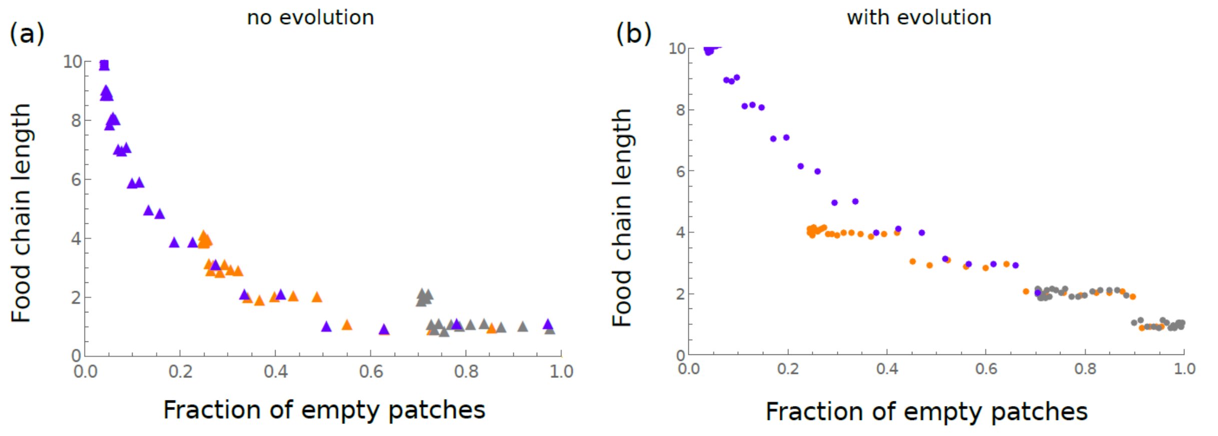

**Supp. Fig. S2** – The association between food-chain length and overall occupancy under varying perturbation rates $\mu$. The relationship between FCL (y-axis) and the proportion of empty patches (x-axis) is shown without evolution (**a**) and with evolution (**b**). Each dot represents a different parameter value, where the perturbation rate $\mu$ was varied (in the same range as *e* in Fig. 5a,b), under three different values of $\psi_0$. In all panels, the three values of $\psi_0$ were -1 (gray dots), -0.1 (orange dots) and -0.01 (blue dots).
